# Onsager regression characterizes living systems in passive measurements

**DOI:** 10.1101/2022.05.15.491928

**Authors:** Till M. Muenker, Gabriel Knotz, Matthias Krüger, Timo Betz

## Abstract

Understanding life is arguably among the most complex scientific problems faced in modern research. From a physics perspective, living systems are complex dynamic entities that operate far from thermodynamic equilibrium.^1–3^ This active, non-equilibrium behaviour, with its constant hunger for energy, allows life to overcome the ever dispersing forces of entropy, and drives cellular organisation.^4, 5^ Unfortunately, most analysis methods provided by the toolbox of statistical mechanics cannot be used in such non-equilibrium situations, forcing researchers to use sophisticated and often invasive approaches to study the mechanistic processes inside living organisms. Here we introduce a new observable coined the mean back relaxation. Based on three-point probabilities, and exploiting Onsager’s regression hypothesis, it extracts additional information from passively observed trajectories compared to classical observables such as the mean squared displacement. We mathematically prove that the mean back relaxation is able to detect broken detailed balance in systems confined in stationary or actively diffusing potentials. We show in experiment and theory that it gives access to the non-equilibrium generating energy and the viscoelastic material properties of a well controlled artificial system, and we experimentally demonstrate that it does so even for a variety of living systems, revealing an astonishing relation between the mean back relaxation and the active mechanical energy. Based on these findings, we conclude that it acts as a new marker of non-equilibrium dynamics. Combining, in a next step, passive fluctuations with the extracted active energy allows to overcome a fundamental barrier in the study of living systems; it gives access to the viscoelastic material properties from passive measurements.

## Main

Any quantitative description of living cells requires detailed knowledge about its mechanical processes, ranging from viscous and elastic local properties to the competition between thermal and active forces. The latter continuously drive any intracellular object, be it an ion, a protein, DNA or even the nucleus. The complex and often randomized nature of active forces acting in living cells makes it hard to dissect active from passive, i.e., thermal components. Such discrimination was up to now only achieved in situations where activity can be tuned, or where the degrees of freedom that break detailed balance can be directly observed, as e.g., in cilia bending modes or by time irreversibility.^6–8^ Besides such special situations, the main experimental method to gain the active forces and the viscoelastic material properties deploys an independent measurement of both, the free particle fluctuations and the mechanical response function representing the viscoelastic material properties. While the former is obtained by non-invasive observation of the erratic particle motion, access to the mechanical properties requires an active force application and the observation of particle motion in response. Comparing the free particle motion with the expected thermal motion predicted by the fluctuation dissipation theorem (FDT),^1, 2, 9–20^ allows then detecting broken detailed balance and the quantification of non-equilibrium contributions, e.g., by defining effective energies. Such direct response measurements, that quantify also the viscoelastic mechanical modulus, were done successfully by optical or magnetic tweezers, AFM and even acoustic forces.^3, 21–23^ While this approach is well accepted, it suffers from several serious problems. The requirement for a simultaneous measurement of nm precise particle motion paired with active and controlled force application in the pN range demands highly specialized equipment that is presently only accessible for few labs worldwide. Additionally, applying an external force, often acting on in-cooperated external particles, inherently disturbs the system, requiring a stringent line of control measurements and still hampers the explanatory power of the obtained results. We present here a new, passive approach that can in principle also be adapted to classical microscopy methods.

### Mean back relaxation detects detailed balance breakdown

Onsager, in his famous regression hypothesis pointed out that the response function, instead of by applying an external force, may equally be measured in response to a *random* force, a statement which is at the heart of the FDT.^24^ However, in living systems, FDT does not hold, and it is difficult to apply this insight. Here we overcome this limitation by introducing a mean back relaxation (MBR) function. In contrast to the mean squared displacement (MSD) which enters the FDT, this function measures the mean particle displacement under the condition that the particle travelled the displacement *d* in the past time interval of length *τ*. In the spirit of Onsager, preselecting displacements *d* plays a similar role as application of a disturbing force (a statement proven for simple models in Material and Methods M 2.2) and the MBR thus quantifies the response to it. More technically, the MBR relates particle positions at three different times, and thus contains more information than the MSD that relates positions at two times.

Before we provide the mathematical definition of the MBR we will illustrate its concept for an overdamped particle confined in a harmonic potential (Fig. 1a). As mentioned, the MBR is defined as the average trajectory of a particle after a recent motion. For example, a particle that moved in the past time *τ* from the potential minimum at position 0 to the position *−d*, is expected to relax on average back to 0 (Fig. 1a upper). Another case that also fulfills the condition is a particle that started at position *−d* and moved to the center in time *τ* (Fig. 1a lower). Fluctuating around the center, this second particle will (on average) contribute zero to MBR. Hence, the here introduced mean back relaxation depends on start and endpoints and the according probabilities. In equilibrium, detailed balance ensures that the probability of moving from *a′* to *b′* (Fig. 1a upper) is exactly the same as from *b′* to *a′* (Fig. 1a lower), i.e., *W*_2_(*a′ → b′*) = *W*_2_(*b′ → a′*). Please note that *W*_2_(*a′ → b′*) is the product of finding the particle at *a′* multiplied by the probability of jumping to *b′*. But only in the first case, the particle will relax back by traveling a displacement *d*, hence averaging over both situations gives a mean relaxation of *d/*2. As we can group all jumps within the potential into such pairs with equal probability but different relaxation path, the mean back relaxation tends to a position that corresponds to exactly half the initial jump displacement, as long as detailed balance is given (see Material and Methods M 2.1).

**Figure 1.**
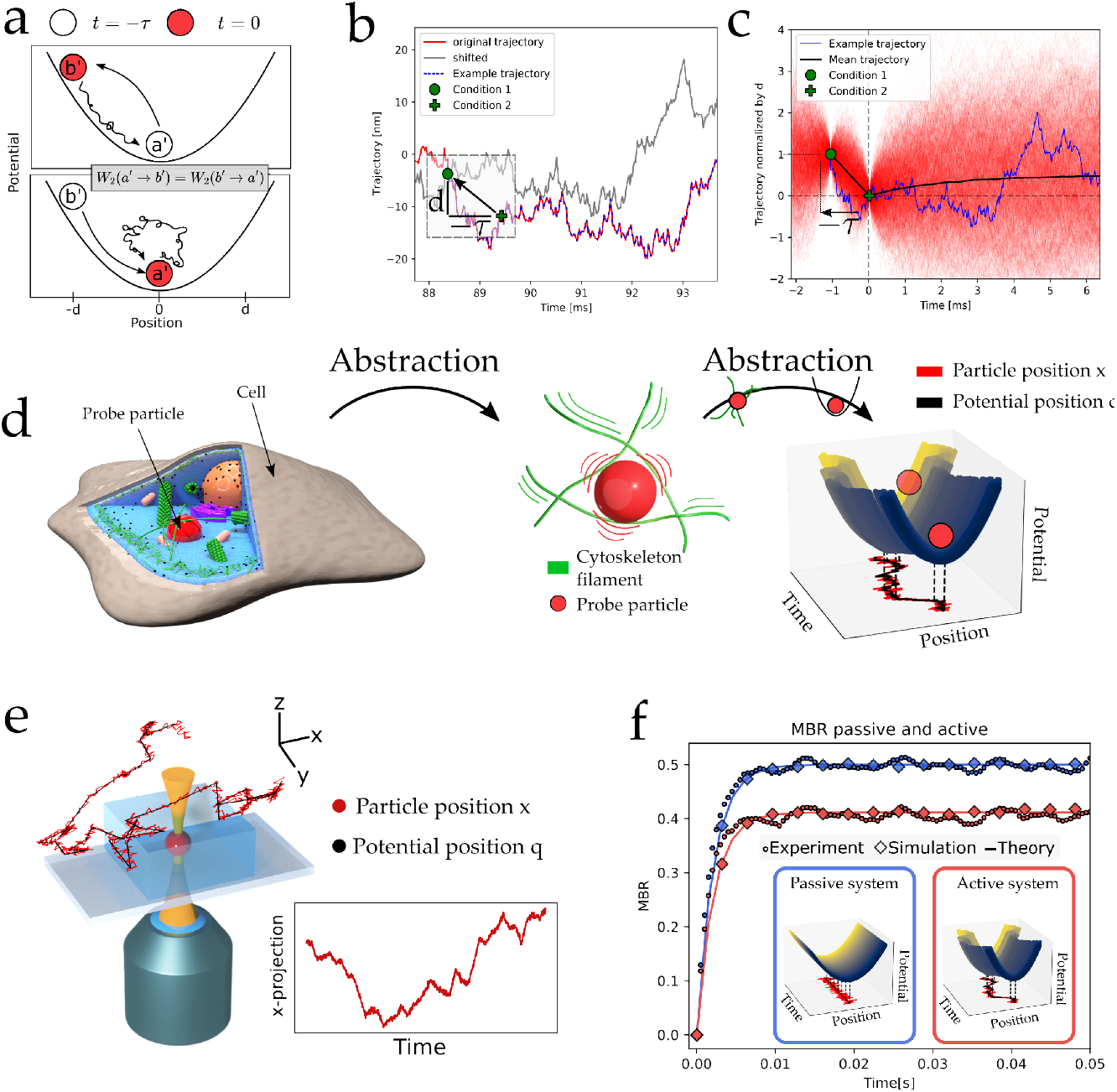
The mean back relaxation (MBR) quantifies a correlation at three different times. a) Schematic of a particle in a harmonic potential. When the particle has moved from the minimum position to an excited state in time *τ*, it will on average go back to the minimum position. Contrary, when it moved from excited state to the minimum, it will on average stay there in the future. b) The MBR relies on particle trajectories that fulfill the condition of a displacement *d* in the history *τ*. This can be visualized by shifting the original trajectory (red) by these parameters (gray). The cross-points mark then trajectories that obey both conditions. c) The average (black) of many (here 10 000, red) such reconditioned trajectories defines the MBR. In blue is the example trajectory from panel a. d) Study of complex cellular material is abstracted by active motion of a harmonic potential.e) Experimental access to the MBR can be achieved via an optical tweezers where the trap is moved to perform a random walk with diffusion coefficient *D*_*q*_. The particle trajectory in the moving trap is obtained in the lab frame using a special detection laser (not shown). f) The resulting MBR for a passive (*D*_*q*_ = 0) and actively driven particles (*D*_*q*_ *>* 0) shows excellent agreement between the theory, the simulations and the experiment.

Example particle trajectories moving in such a harmonic potential can be conveniently obtained using Brownian dynamics simulations (Fig. 1b). The trajectories fulfilling our precondition can be visually selected by shifting a copy of the trajectory by *d* and *−τ* (Fig. 1b, gray). The crossing points of original and shifted trajectories are collected and overlaid as initial positions for the MBR curves. As there are few crossing points (Fig. 1b), we increase statistics by using any value of *d* (keeping *τ* fixed), and then overlaying the resulting displacements normalized by *d*. Fig. 1c shows an overlay of 10,000 such trajectories (light red) and the resulting mean (black). The latter shows a kink at time *t* = 0, implying that the particle on average *reverts* the conditioned displacement. As introduced above, this is intuitive as the conditioning selects particles that just moved up in the potential, and thus, for positive times on average relax back down. This intuition may explain the sign of the MBR, but not its magnitude. The particle travels back *half* of its initial drift. This nontrivial ratio of 1*/*2 can be derived analytically using probability distributions, and it holds generally for a particle in a stationary potential, for any dynamics obeying detailed balance (Material and Methods M 2.1). In summary, the mean traveled distance as measured in the MBR is finite because of the preselection condition, and the reason for its sign is the average (back) relaxation of the particle to the potential minimum. Formally, the MBR is defined via the three point joint probability *W*_3_(*x, t*; *x′, t′*; *x″, t″*) of finding the particle at *x* at time *t* and at *x′* and *x″* at the earlier times *t′* and *t″*(Material and Methods M 1),

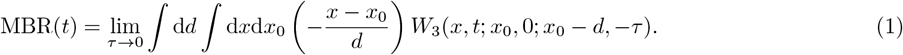

The MBR depends on *W*_3_, and we expect it to contain more information than the MSD, which involves two times only.

Although aiming at understanding the complexity of a living cell (Fig. 1d) we first simplified our analysis to a situation where a particle rests in a local viscoelastic cage, such as a cytoskeletal network. This cage presents a local trap for the particle that is then modeled by a harmonic potential, centered at position *q*(*t*). Hence, we use the harmonic potential to model the local trapping by the cytoskeleton (Fig. 1d). As introduced above, the particle at position *x* performs a random motion with a diffusion constant *D*. The active forces as found in living cells can distort the cytoskeletal background (Fig 1d, center), mimicked by moving the harmonic potential also on a random trajectory (Fig. 1d, right) with diffusion constant *D*_*q*_, *unaffected* by the particle. Since the movement of the trap *q*(*t*) is predefined, it violates the Einstein relation so that the trap diffusion is neither coupled to the particle nor to thermal energy. While in the introductory example above (Fig. 1a), the harmonic potential is at rest, we now introduce with *q* a second degree of freedom, namely the trap position. This system of two coupled degrees of freedom (particle and trap position) violates detailed balance because the trap moves independently of the particle – dragging it like a (random) horse drags a cart. If *x* and *q* are experimentally accessible, braking of detailed balance can be detected using classical approaches such as comparing the motion of particle and trap with their time reversal. This gives access to time antisymmetric quantities such as entropy production.^25^ To mimic an experiment in a living cell where we do not know the distortion of the cytoskeletal background, we pretend to be blind to the trap position, i.e., treat *q* as a *hidden* degree of freedom. When studying complex systems, such presence of hidden degrees is the typical case. Then the entropy production cannot easily be measured and detecting non-equilibrium is nontrivial.

What can we learn for this system from conditioned trajectories? Spontaneous fluctuations push the particle to exited states in the potential, from where it will (on average) relax to the potential minimum, *x* = *q*. The MBR thus equals the normalized mean distance between *x* and *q* at time *t* = 0, 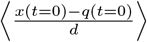, under conditioning. Thereby the MBR provides information about the potential position *q*. This is remarkable, as we thus obtain information about the hidden degree of freedom *q*, despite being experimentally blind to it. More precisely, as the value of MBR depends on *q*(*t* = 0) (see above), it tests the joint distribution of *q* and *x*, and a deviation from 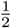 can only occur if detailed balance for *q* and *x* is broken (Eq. (16) in M 2.2).

Explicitly, for the given model of particle and diffusing trap, the MBR is found analytically (Material and Methods M 2.2)

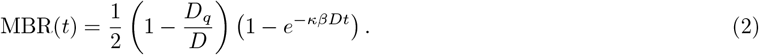

Here *κ* is the spring stiffness of the harmonic potential and *β* = (*k*_*B*_*T*)^*−*1^ is inverse thermal energy. As noted, for large time *t*, the MBR equals 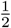 for the equilibrium case, which corresponds to the situation where the harmonic potential is stationary. This is mathematically equivalent to a situation with a trap diffusion constant of *D*_*q*_ = 0, keeping mind that we can experimentally control *D*_*q*_. For non-equilibrium cases, i.e., for finite *D*_*q*_, detailed balance is broken, and the long time MBR takes smaller values (M 2.2): For large *D*_*q*_, the particle is strongly dragged by the potential, so that the displacements in time span *τ* and afterwards are typically directed in the *same* direction, i.e., there is a *forward* relaxation, and the MBR is negative.

### Experimental validation of the MBR’s predictive power

The experimental realization of the introduced theoretical model was achieved by trapping a 1*μ*m polystyrene particle, suspended in a Newtonian fluid, with a movable optical tweezers (Fig. 1e, Material and Methods M 4). The particle motion was measured using an additional, fixed, low power detection laser with nm and *μs* resolution. Implementing a random trap trajectory with a diffusion constant *D*_*q*_ enabled us to directly confirm the theory and simulations of the MBR (Fig. 1f, 2a). The MBR was calculated according to SI 1. As predicted, the experimental realization of a passive system with *D*_*q*_ = 0 showed a rapid relaxation of the MBR to the expected value of 1/2. But randomly moving the trap by increasing *D*_*q*_ *>* 0, and hence driving the system to a non-equilibrium state, led to a decrease of the MBR and its long time limit (Fig. 2b). To verify experimentally the Onsager regression hypothesis, we checked if the MBR reflects the mechanical properties of the studied system. Indeed, by fitting the time dependent relaxation of the MBR to the theoretical expression we obtained the relative non-equilibrium contribution represented by the trap diffusion *D*_*q*_ and the mechanical properties of the experimental system represented by the stiffness of the optical trap *κ* and the friction coefficient *γ* (Fig. 2a,b). A surge in active driving by an increase of the trap motion confirmed that the MBR can become negative as suggested by Eq. (2) (Fig. 2b). One striking prediction of the MBR is that already its long time limit can be used as a simple way to quantify the active non-equilibrium driving. Again, we find an excellent agreement between the applied active motion, quantified by *D*_*q*_*/D* and the expected MBR value (Fig. 2c), where negative values of the MBR implicate that the motion is dominated by the trap motion, and positive values mark a temperature dominated motion. These results suggest that the MBR allows determining if the system is out of equilibrium, and at the same time quantifying the relative intensity of the active motion (here *D*_*q*_). Hence, we have demonstrated experimental access to the recently introduced active diffusion^26–29^ by a simple and noninvasive trajectory analysis.

**Figure 2.**
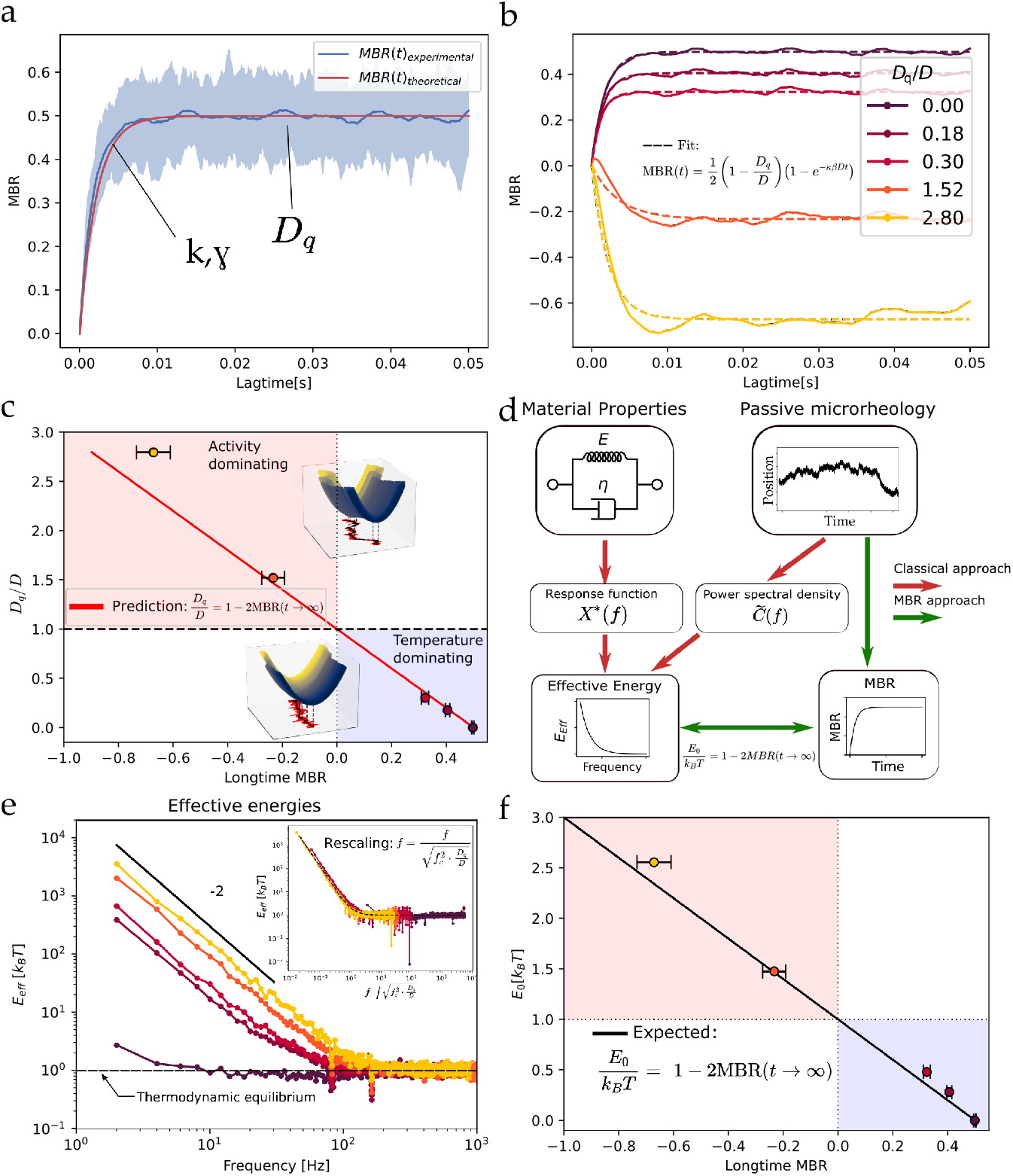
MBR gives access to mechanics and activity. a) Fitting the theoretical model to the MBR data allow extracting the mechanical properties and activity as quantified by the active trap diffusion. b) Increasing the active trap diffusion leads to changes in the long term limit of the MBR. c) Already the long term limit of the MBR allows inferring the relative active diffusion, where a negative value is identified with activity dominating and a positive value means the fluctuations are temperature dominated. d) The effective energy is a common quantity used to determine activity. This can be inferred from measurements of mechanics and spontaneous fluctuations, but also from the MBR. e) Effective energy of the movable trap for different activities *D*_*q*_. Inset: Using the theoretical prediction, effective energies can be rescaled to master curve using known values of *f*_*c*_, *D* and *D*_*q*_. f) The effective energy can be determined from the long time limit of the MBR, similar to the active diffusion

### Measure of activity from passive observation

Considering this important new access to non-equilibrium properties, we wondered if the long time limit of the MBR can also be used to determine another key quantity for a thermodynamic description of active systems, namely the energy that is injected into and dissipated by the system. We quantify it as *effective energy E*_eff_ (*f*) that is defined by comparing the particle fluctuations for a frequency *f* with the Brownian motion expected by FDT (Fig. 2d red path). Excess fluctuations exceeding the Brownian level are attributed to the active processes as illustrated in Fig. 2e (see SI 4 for a definition).^1, 3, 30^ In our system of a randomly moving trap, both, experimentally as well as analytically (SI 4), the effective energy follows 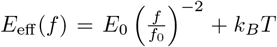 with the frequency *f*_0_ = (*κD*)*/*(2*πk*_*B*_*T*) in this model. The analytical divergence at *f →* 0, hence infinite time, is experimentally not reached because we only record traces for 1 second. Astonishingly, the analytical analysis predicts a direct relation between *E*_0_ and the MBR,

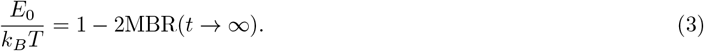

It is in perfect agreement with the data (Fig. 2f). Hence, the larger distance between the MBR from 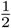, the larger the amplitude *E*_0_ of the effective energy. While (3) can be extended to include several particles or memory, the validity of this relation in other systems and models is unknown. We continue based on the hypothesis that the MBR gives access to dissipated energies in other non-equilibrium systems as well (Fig. 2d green).

Analyzing a system with broken detailed balance, we wondered whether additional information may be gained from *time reversed* particle trajectories. However, in this model, the reversed trajectories are statistically *indistinguishable* from the original ones (Material and Methods M 2.2, Extended data Fig.5). This means, e.g., that MBRs evaluated from reversed or original trajectories are identical, despite detailed balance being broken.

### MBR gives access to non-equilibrium energy in cells

To test if the MBR can also be used in more complex and realistic environments, we turned to analyzing the erratic motion of particles embedded in living cells, which represents a complicated and highly relevant system. As active forces generated by molecular motors provide the dominant contribution to intracellular particle motion, this system is far from equilibrium. A purely passive measurement of the particle trajectory is thus not expected to provide sufficient information about the mechanical properties or the active contributions, as detailed above.^1, 2, 31^ However, with respect to our findings that the MBR, carrying additional information compared to the MSD, may open the door to quantify active non-equilibrium motion we decided to test its potential in the study of cellular mechanics and activity. To this end, we allowed cervical cancer cells (HeLa) to phagocytoze 1*μ*m diameter polystyrene particles which we then used as proxies for intracellular organelles, such as lysosomes as done previously.^3^ Using the optical tweezers setup (Fig. 3a, Material and Methods M 3), we can experimentally determine both the spontaneous fluctuation by pure passive observation and then directly access the viscoelastic properties in terms of a response function *χ*(*f*) (Fig. 3b) and the viscoelastic shear modulus *G*\*(*f*) (Fig. 3c) using active microrheology (AR) (Material and Methods M 5). We have hence direct experimental access to the effective energy (Material and Methods: Eq. (36)) and the mechanical properties which brings us in a perfect position to test the potential of the MBR. The main aim here is to extend a passive microrheology approach to active systems, which has not been possible before.

**Figure 3.**
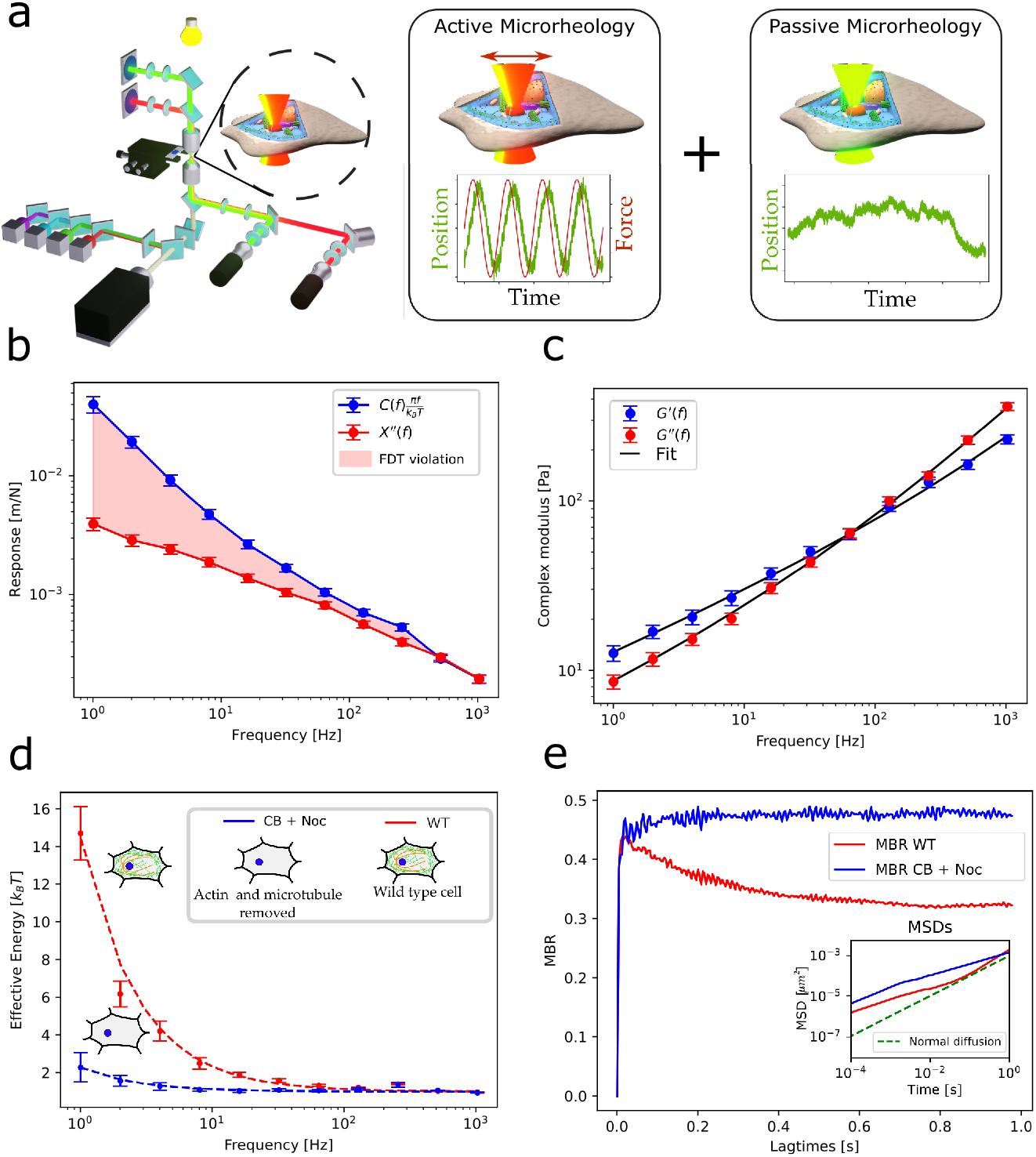
MBR as new quantity to analyse living cells: a) combining a fluorescent microscope with optical tweezers and detection lasers allows simultaneous measurement of the viscoelastic properties, the effective energy and the MBR in living cells. b) Comparing the response expected from the fluctuations and the direct measurements shows violation of the fluctuation dissipation theorem. c) Direct measurement of the response function gives access to the complex shear modulus. d) Comparing the effective energy in WT cells (red) and cells with pharmacologically reduced motor activity (blue) shows a breakdown of effective energy in the drugged, and hence passive cells. e) the MBR of passivated cells (blue) gives similar results as a passive, optically trapped particle. In active WT cells, activity is indicated by the deviation of the MBR long term limit from 1/2. Inset) The MSD of the two curves shows superdiffusive behaviour in the WT cells, which is not the case in the passive cells.

To study the MBR in active and passive cellular systems, we compared the effective energy and the MBR of untreated WT cells with passivated cells. Here the active forces are drastically reduced by the disruption of the actin (Cytochalasin B, CB) and microtubule (Nocodazole, Noc) cytoskeleton on which the molecular motor proteins apply their forces (Material and Methods M 8). As expected, the statistics of particle motions in this depleted cell is similar to an equilibrium system, i.e., the effective energy remains close to *k*_*B*_*T*, while in an intact cell, *E*_eff_ can be much larger than *k*_*B*_*T* for low frequencies (Fig. 3b).

Turning to the MBR (SI 1) now shows the potential power of this new quantity. In the depleted, but still living cells, we found that the predicted long time limit of 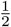 is indeed closely approached, demonstrating that this general behaviour also holds for this complex passive system. Turning to the MBR of active living cells, we first see an overall different shape of the MBR, as an initial steep increase follows a decay to the long time limit. As seen above for the model system, the MBR converges in the active case to a long time limit smaller than 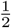. The activity can also be confirmed by investigating the MSD (Fig. 3e inset), as in the active cell, the particle motion becomes superdiffusive in the long time limit. This supports the above hypothesis that, similarly to the actively moving trap system, a deviation of the MBR from the equilibrium value 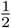 can be used to detect active motion.

### MBR allows passive microrheology in active non-equilibrium systems

This raises the question if the MBR can be used to also quantify activity in living cells. To test this, we measured mechanics and activity in terms of effective energy as well as the MBR of 7 different cell types and conditions ranging from epithelial over cancer, kidney, muscle and immune cells (Extended Data Fig 2, Material and Methods M 7). As these different cell types show variable viscoelastic and active properties, this broad approach allowed exploring the relation between MBR, activity and mechanics in more detail. Consistent with previous experiments,^3^ we find that the effective energy in cells can be expressed by a powerlaw, *E*_eff_ = *E*_0_(*f/f*_0_)^*v*^ + *k*_*B*_*T* (Fig. 4a), where the frequency *f*_0_ is set to 1 Hz. While the prefactor *E*_0_, which quantifies the extend of metabolic energy dissipated in the system, varied drastically between different cells, the powerlaw exponent was found to only vary non-significantly around a value of *v ≈ −*1. To infer a potential relation, we inspected the effective energy *E*_0_ as a function of the MBR long time limit as shown in Fig. 4b. Common state of the art expectation is that it is difficult to quantify activity from passive measurements.^1–3^ Again, we calculated the MBR for all cell types according to SI 1. Keeping in mind that the MBR is inferred from such passive observation, to our great surprise, we found an impressively consistent linear relation between the two quantities. This striking dependence opens the door to experimentally access not only the effective energy that is driving intracellular objects but also to determine the viscoelastic properties inside a cell by simply observing a particle trajectory.

**Figure 4.**
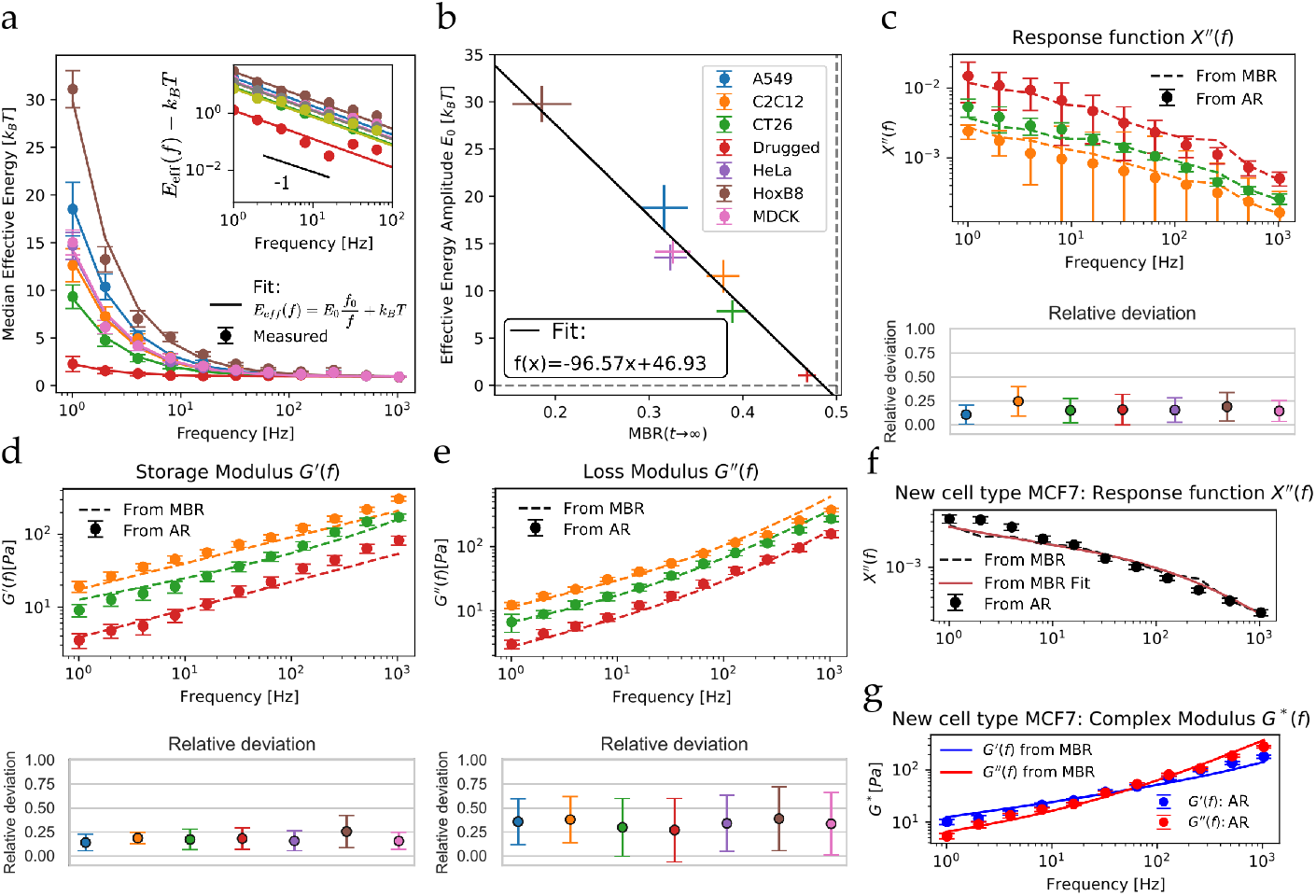
MBR long term limit relates to effective energy in cells via a linear mastercurve. a) Analysing the effective energy of 7 different cellular situations shows powerlaw behaviour. Inset) Detailed analysis shows a general powerlaw exponent of -1. b) Plotting the prefactor of the effective energy *E*_0_ against the long time limit of the MBR (at 1s) shows a linear dependence, where the values for all cells are found on a mastercurve. c) Using the FDT with the correction for the effective energy and the *E*_0_ from the mastercurve allows extracting the dissipative response function purely from passive measurements of the MBR and the power spectral density. The error between the direct measurements and the inference via the MBR is typically smaller than 10%. d,e) Fitting a generally accepted phenomenological model to the dissipative response function allows extracting the real and imaginary part of the shear modulus. Again, the error between the direct measurements and the values inferred from the passive measurements is typically below 16%. f) Applying the method to a new, yet unused cell confirms that the MBR can be used to extract the dissipative response function from a passive measurement of an active system. g) The resulting shear modulus is in excellent agreement with the direct measurements.

Exploiting these findings, we define an adapted version of the fluctuation dissipation theorem for active systems. This is based on combining the spontaneous fluctuations as quantified by the powerspectral density *C*(*f*) and the analytical expression for the effective energy that are related to the dissipative response function *χ″*(*f*) (Fig. 4c) via (Material and Methods M 5):

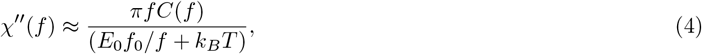

When now comparing the response function that is predicted using *E*_0_ obtained via the MBR (Fig. 4b) with the independently measured response function via active microrheology (AR) (Fig. 4c upper), we found an excellent agreement with an average relative error of 16% (Fig. 4c lower). Using this approach we can successfully solve one of the most hindering problems of modern mechanical biology, as we determine both, the active mechanical energy dissipated, and the mechanical properties defined by the response function of a living cellular system by purely observing the passive particle motion. This approach is fundamentally different to classical passive microrheology which can only be used in thermodynamic equilibrium and would yield an average relative error of 236% in this situation (Extended Data Fig. 3). Once knowing the dissipation of the system we can either exploit Kramers-Kronig relation, or a typical phenomenological model such as a coupled springpot to directly gain the real and complex part of the viscoelastic shear modulus as shown in (Fig. 4d,e, Material and Methods M 6).^3, 23, 32^ To test the predictive power of our approach, we used the here developed new formalism to analyze a new cell type (MCF7), that was not previously included in the analysis. Impressively, we can compute with very good agreement the independently measured imaginary response function (Fig. 4f, error 16.3%) and the complex shear modulus (Fig. 4g, error G’: 16.8%, error G”: 15,2%) by only using the passive particle trajectory and the phenomenological laws established above.

## Conclusion

We demonstrate that the new quantity of mean back relaxation allows new insights into non-equilibrium systems, which cannot be given by other known analysis. Our analysis demonstrates the predictive power of the MBR by re-establishing passive microrheology to analyse active cellular systems, which now makes a rapid and detailed local and temporal characterization of the viscoelastic material properties inside cells and active biological system possible. Compared to the current state of the art (Fig. 5), we do not require a combined analysis of active and passive measurements. Furthermore, the required temporal and spatial resolution of the particle trajectories can in principle be obtained in standard research microscopes equipped with high speed CMOS cameras. Given sufficient illumination and using standard tracking algorithms, subpixel resolution at frame rates of multiple kilohertz can be achieved.^33–35^ This allows to use the MBR approach in most modern research labs. The simple access has large implications for new models and new theoretical approaches to study one of the most important scientific subjects of our time, represented by actively driven microscopic systems.

**Figure 5.**
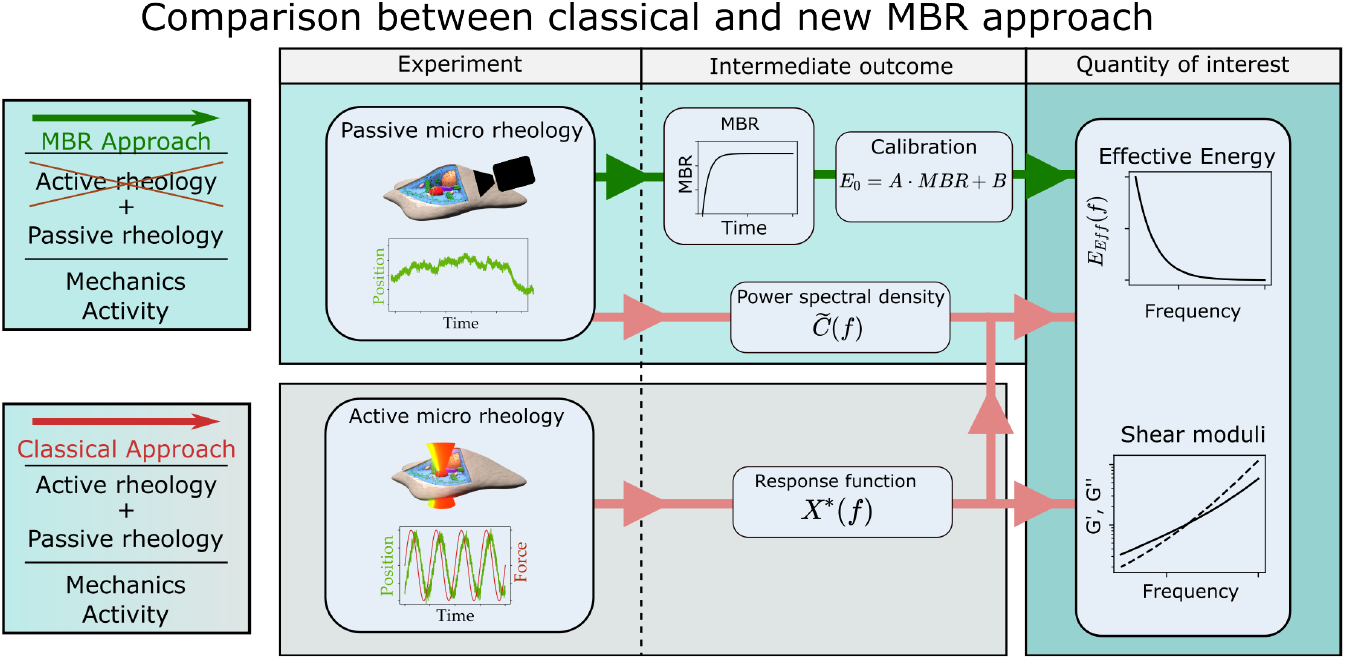
Comparison between the classical approach which relies on a simultaneous access to the freely fluctuating particle trajectories and the optical tweezers based rheology. In contrast the access via the MBR only requires the passive observation of the particle trajectory, and yield eventually the same results as the classical approach.

Correlating particle positions measured at three different times, like in the MBR, is expected to distinguish equilibrium from nonequilibrium trajectories, as has been shown theoretically for a discrete model.^36^ But such idea has not been put forward for continuous (experimental) systems. The MBR introduced here can not only distinguish equilibrium from nonequilibrium systems, it also *quantifies* the distance from equilibrium. As detailed in Eq. (20)(Material and Methods M 2.2) for a particle in a potential, conditioning trajectories for MBR yields a *shifted* particle distribution and has the same effect as application of a short force pulse. Recalling Onsager, the selected trajectories are thus equivalent to a perturbed initial state, and the MBR measures its relaxation. Because this perturbation is given by the ratio of active and thermal energies (Eq. (20)), the MBR allows to quantify active energies from passive measurements. These statements are proven analytically in paradigmatic active models and experimentally observed in a system as complex as a living cell. Future theoretical work can investigate the relation to, e.g., first passage times and higher order moments,^37–39^ the concept of biased ensembles,^40^ large deviation theory^41, 42^ and entropy production.^43–47^

This opens the door for a new experimental and theoretical access to quantify life from a mechanical point of view.

## Material and Methods

### M 1 Mean back relaxation: Definition

The mean squared displacement cannot detect the breaking of detailed balance.^36, 48^ It is based on the joint probability *W*_2_(*x, t*; *x′, t′*) of particle position *x* at time *t* and the earlier position *x′* at time *t′*. Here we aim to explore an observable based on the *three* point probability. To illustrate this consider the mean traveled distance after selecting preconditioned trajectories. A simple and intuitive condition is picking trajectories which traveled a displacement *d* in the immediate history, i.e., between times *−τ* and zero. We define the *mean back relaxation* (MBR) via

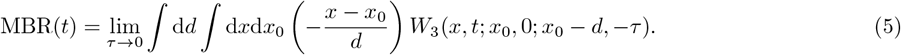

*W*_3_(*x, t*; *x′, t′*; *x″, t″*) is the joint probability of finding the particle at positions *x, x′* and *x″* at the specified times, with *t* and *τ* positive in Eq. (5). For simplicity of discussion we take the limit *τ →* 0 (only after evaluation of the integral). In a trap the particle will often move in a direction opposite to *d*, because it has to travel back to the center of the trap. So we multiply with a minus sign to obtain a positive result for the MBR in equilibrium. In simple cases the displacement for positive times is linear in *d* (see comments below) and we normalize by *d*. When determining the MBR from trajectories, it is useful to introduce a small lower cutoff for *d* to avoid division by small numbers. After integration over displacements *d*, the MBR defined in Eq. (5) is a function of correlation time *t* only.

### M 2 Exact relations for the MBR

#### M 2.1 MBR is a non-equilibrium marker – bound case

We start by consider the long time limit of the MBR in a stationary system, i.e., in a system with a stationary distribution *P*_stat_(*x*) with finite mean. We can then decompose the three point probability into 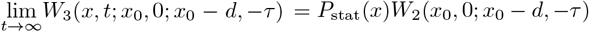, assuming finite correlation times. Inserting in Eq. (5) yields

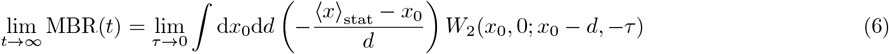

where ⟨*x*⟩_stat_ = ∫d*xP*_stat_(*x*)*x* the mean of *x* with respect to the stationary distribution. We split the integral in two parts, substituting in one of them *x*_0_ *→ x*_0_ *− d* and *d →* (*−d*). This yields an exact reformulation of Eq. (6)

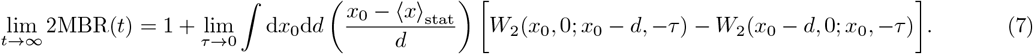

Notably, the difference *W*_2_(*x*_0_, 0; *x*_0_*−d, −τ*)*−W*_2_(*x*_0_*−d*, 0; *x*_0_, *−τ*) enters. With detailed balance fulfilled, i.e., *W*_2_(*x′, t′*; *x, t*) = *W*_2_(*x, t′*; *x′, t*), the term in the integral vanishes, and

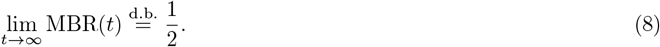

This states that, if the long time value of the MBR deviates from 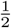, detailed balance must be broken. MBR thus detects deviation from detailed balance, i.e., deviation from equilibrium. In deriving Eq. (8), we have not made any assumptions regarding the dynamics. The given statements thus hold also, e.g., in cases with inertia, memory, or interactions, including non-Markovianity.

It is illustrative to reformulate this statement in terms of stochastic entropy production for trajectories *x*(*t*) with probability *P* as

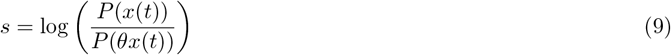

with the time reversal operator *θx*(*t*) = *x*(*τ − t*). For the considered steady state scenario, the expression Eq. (7) is equivalent to the fluctuation theorem

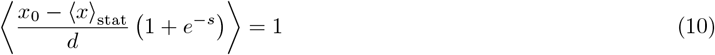

which relates the long time MBR and the stochastic entropy production.

As a specific example we regard a Brownian particle in a harmonic potential 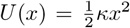 with diffusion *D* and temperature 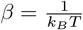. Because this system is Markovian, the three point probability can be expressed as

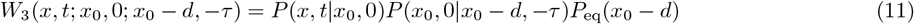

with the conditional probability 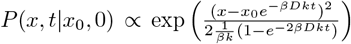, which can be found by using Gaussianity and solving the corresponding Langevin equation. By introducing the displacement *b* = *x − x*_0_ and completing the square to eliminate *x*_0_ the MBR is reduced to

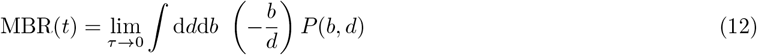

with 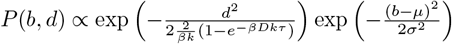, with mean and variance

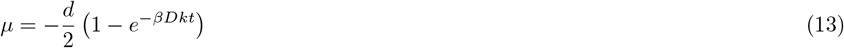

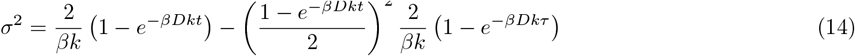

Note that in this example *μ* is independent of *τ* which makes the limit *τ →* 0 trivial after the remaining Gaussian integrations have been performed. Thus the MBR of this system is given by

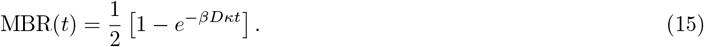

It is positive for all times *t* and approaches the long time value of 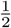 as expected. In contrast, for finite time, the relaxation is governed by system details such as *βDκ*.

#### M 2.2 MBR is non-equilibrium marker – active potential model

To create a simple non-equilibrium system that mimics a changing background, we extend the above to a particle (position *x*) subject to a potential, whose position *q* (e.g. its minimum) performs a random walk with diffusion coefficient *D*_*q*_. This is an active system, because *q* is assumed not to be influenced by *x*, as described in the main text. *q* thus may mimic an active motor, which is insensitive to the particle at *x*. In steady state, the distance *x – q* takes a time independent mean value ⟨*x − q*⟩_stat_. The steps leading to Eq. (7) can be repeated to arrive at the *t → ∞* limit of MBR

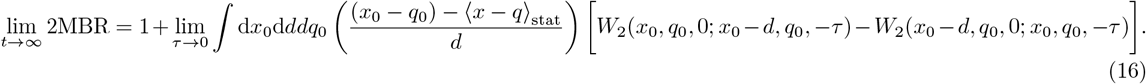

Eq. (16) shows that, also in this model, a deviation of MBR from 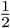 can only occur if detailed balance is broken. Again, for the derivation of Eq. (16) we made no assumptions about the dynamics of the particle *x*.

We turn to the specific example of a Brownian particle in a potential *U* (*x*), diffusing with *D*_*q*_, as discussed in the main text. The Langevin equations of this system are

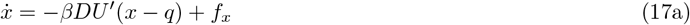

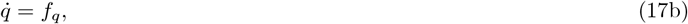

with the independent white noise forces *f*_*x*_ and *f*_*q*_ given by ⟨*f*_*x*_(*t*)*f*_*x*_(*t′*)⟩ = 2*Dδ*(*t − t′*) and ⟨*f*_*q*_(*t*)*f*_*q*_(*t′*)⟩ = 2*D*_*q*_*δ*(*t − t′*). It is useful to focus on the relative coordinate *x – q* to obtain the stationary distribution of the system,

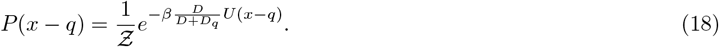

This system thus follows a Boltzmann distribution, however, with an effective temperature 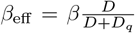. The short time dynamics of the *x* variable is governed by its diffusion constant *D*. Therefore, for small *τ*, the dynamics of the system is diffusive, added by a drift due to the potential gradient.^49^ The short time evolution of the joint probability for a travelled displacement *d* is thus given by^49^

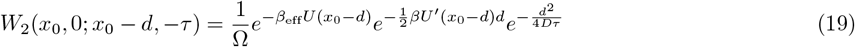

with normalization Ω. For small *d* and *τ* we can interpret the expression in the exponent as a Taylor expansion in *d* around *x*_0_ *− d*. Using the standard chain rule from probability calculus for the conditioned probability, 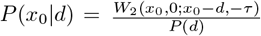, with *P* (*d*) = ∫ *dx*_0_*W*_2_(*x*_0_, 0; *x*_0_ *− d, −τ*),

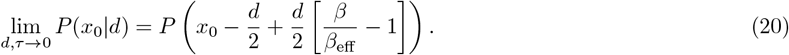

The conditioned distribution therefore corresponds to the original distribution, however shifted by a certain amount. Notably, this distribution is exactly equal to the one obtained when applying a force pulse of length *τ* and magnitude 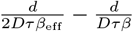, i.e., directly related to the effective energy.

For long times *t*, we have

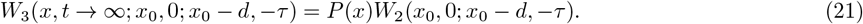

Using this in Eq. (5) yields a long time MBR of ^1^

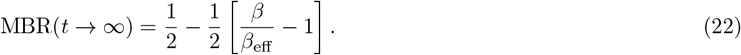

In equilibrium with *β* = *β*_eff_ the term in the brackets vanishes and we obtain 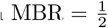 as expected. In this particular model deviations of the equilibrium value are directly related to the ration of *β* and *β*_eff_. It should be noted that such analysis using effective temperature (as also discussed in the main text) requires that the system shows a Boltzmann distribution at long times.

In the main text we discuss the simplest case 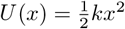. The analytic solution of this model can be found as

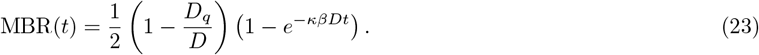

This relation is discussed and verified in the main text. Notably, the finite diffusivity of *q* is predicted to change only the overall prefactor of the MBR, however in a drastic manner: In contrast to the equilibrium system at *D*_*q*_ = 0, it deviates from 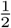 and can take *negative* values. This equation shows that the deviation of the long time value of the MBR is directly related to the deviation of *β*_eff_ from *β*. Using, as in the main text, 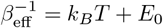, we obtain

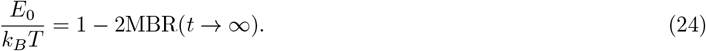

A remark on measuring time reversal symmetry for the diffusing potential. If *x* and *q* are (experimentally) accessible, an obvious measure is the entropy production of the system, i.e., the work of the motor particle 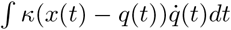. However, when only the particle position *x* is known, entropy production cannot be measured. Furthermore, it is insightful to regard the time reversed dynamics of the motor model,

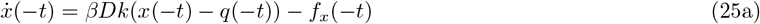

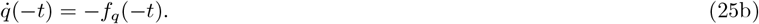

By formally solving the forward dynamics *x*_*f*_ in Eq. (17) and comparing to the solution of the backward dynamics *x*_*b*_ via Eq. (25) we get

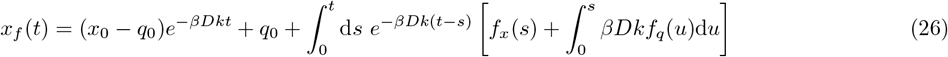

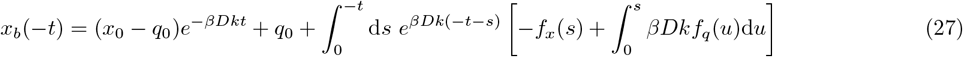

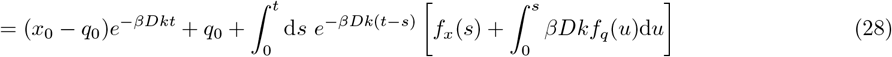

In the last step we mapped *s → −s* and *u → −u*, using *f*_*x*_(*−s*) = *f*_*x*_(*s*) and *f*_*q*_(*−s*) = *f*_*q*_(*s*). So we find that the dynamics in forward and backward time-direction of the dynamics of the particle are *statistically identical*. This proves the statement given in the main text, that the MBR evaluated from backward trajectories is identical to the one from forward trajectories for the diffusing potential.

### M 3 Optical tweezers setup

The optical tweezers setup (Extended Data Fig 4) is based on 2 diode lasers which are used for trapping and position detection of the particles respectively. An 808 nm, 250 mW laser (LU0808M250, 808 nm, 250 mW Lumics GmbH, Berlin, Germany), ensuring minimal laser absorption in biological tissue,^50^ is used to trap the particles and for measurement of the forces exerted to these particles. The collimated laser beam is expanded by a 5x telescope setup to slightly overfill the back aperture of the objective. The trapping laser can be steered by an extra thin mirror (diameter 12.7 μm x thickness 1 mm custom build, Chroma, Bellows Falls, USA) glued onto a piezo tilting platform (S-331, Physik Instrumente (PI) GmbH & Co. KG, Karlsruhe, Germany). Using a 4f configuration, the plane of the piezo rotation mirror is projected onto the back aperture of the objective, allowing trap positioning while optimal entering the objective.^51^ The high NA objective (CFI Plan Apochromat VC 60XC WI NA 1.2, Nikon) focuses the beam to a diffraction limited spot in the sample plane. Sample position can be manipulated with a piezo stage (MAX311D/M, Thorlabs, New Jersey, USA) of 20 μm fine travel distance. A 1.4 NA condenser (extracted from Deimus T-10i, Impetux, Barcelona, Spain) is ensuring that almost all of the persistent light is collected and can be used for accurate force detection. The back focal plane (BFP) of the condenser is then imaged onto a Position Sensitive Detector (order nr. 5000011, First Sensor, Berlin, Germany), where, given correct alignment, the trapping force can be detected. This approach does not require daily calibration, and the correct force measurement was critically crosschecked using the drag force method. A more detailed description of the force detection method can be found elsewhere.^52^ Position detection was achieved using back focal plane interferometry.^53^ A 976 nm (BL976-PAG500, 976 nm, 500 mW, Thorlabs, New Jersey, USA) IR laser is first expanded by a factor of 3 and then reduced in intensity by a neutral density filter (0.1% transmission, NE530B, Thorlabs, New Jersey, USA). This allows stable laser operation at high diode driver currents, while the laser power at the sample plane is in the order of a few mW. Next, the beam is coupled into the optical path via a dichroic mirror (DM) (DMLP900, Thorlabs, New Jersey, USA). Above the condenser both beams are again separated from the illumination path. A 785 nm notch beam splitter (F78-785, AHF Analysetechnik AG, Tübingen, Germany) is used to reflect the 808 nm trapping laser on the force detector, while letting the illumination light and the 976 nm position detection laser light pass. Then the 976 nm laser is reflected on the force detector using a 650 nm short pass dichroic mirror (DMSP650R, Thorlabs, New Jersey, USA). The BFP of the condenser is imaged onto a Quadrant photo diode (QPD, PDQ80A, Thorlabs, New Jersey, USA) where a signal proportional to the particle displacement can be measured. An aperture is located in front of the QPD in order to allow axial position detection.

Brightfield illumination is achieved using a 620 nm LED (SOLIS-620D, Thorlabs, New Jersey, USA). The imaging beam is decoupled from the beam path using a 775 nm short pass dichroic mirror (ZT775sp-2P, Chroma, Bellows Falls, USA). Using a tube lens (TTL200-UVB, Thorlabs, New Jersey, USA) the object plane is finally projected onto the camera (ORCA FLASH4.0 V3 QE80, Hamamatsu Photonics, Japan).

The setup was further extended for fluorescence microscopy. A 4-Wavelength LED Source (LED4D255: 405/490/565/660 nm, Thorlabs, New Jersey, USA) is coupled into the beam path using a 4 band beam splitter (ZT405/488/561/647rpc, Chroma Technology, Bellows Falls, USA) which reflects the excitation wave length and transmits emission and bright field light.

In order to keep the sample at a constant temperature of 37±C both, objective and condenser can be heated with an objective heating system (H401-T-Controller, Objective Heater, Okolab, Pozzuoli, Italy).

### M 4 Experimental realization of the diffusing potential

To experimentally realize a driven harmonic potential in a Newtonian fluid, we employed the optical tweezers setup described afore. Probe particles (Polybead^®^ Microspheres 1 μm, Polyscience, Inc) were suspended in 60% Glycerol (ROTIPURAN^®^, Roth) at a dilution of 1:10000 and then inserted to the probe chamber. The probe chamber consists of 2 cover slips (22×55×0.15 mm^3^ VWR) glued together with double sided tape (DST1950, Thorlabs, New Jersey, USA) at both ends to create a chamber of 200 μm height. To mimic the harmonic potential, the 808 nm optical tweezers laser was focused onto a probe particle and thereby, for small particle displacements, trapping it in a harmonic potential.

To realize different kinds of activity, a voltage signal representing a discrete random walk was applied to the x-axis of the piezo mirror controlling the position of the trapping laser. The signal *V* (*t*) is given by: *V* (*t*) = *V* (*t −* Δ*t*) + *V*_0_(*σ*_*T*_) where *V*_0_(*σ*_*T*_) is randomly drawn from normal distribution with variance 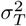. For all experiments, Δ*t* was chosen as 1 ms and *σ*_*T*_ controlled the level of activity. For each experiment, the feedback signal of the mirror was monitored and translated into a laser displacement signal using a previously obtained calibration curve. To determine the trap diffusion coefficient *D*_*q*_ with highest accuracy, the laser displacement signal was used to calculate the mean square displacement (MSD) from which *D*_*q*_ was derived according to MSD(*t*) = 2*D*_*q*_*t*.

During experiments, a second laser was used to monitor the particle position. This laser was operating at a very low power and did not apply any relevant additional forces to the probe particle. In order to measure the trap stiffness *k* of the optical tweezers and the viscosity *η* of the Glycerol dilution, the particle position was monitored while the harmonic potential was stationary. The trap stiffness *k* was obtained from the variance 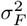 by usage of equipartition theorem 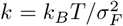. From the short time MSD of the particle position signal, we obtained the diffusion constant *D* and thus also the viscosity *η* via *D* = *k*_*B*_*T/*(6*πηR*) with particle radius R=0.5 μm given.

For each activity level of the trap *D*_*q*_ ∈ {0, 0.18, 0.3, 1.52, 2.8}, 100 different trajectories were recorded, each containing 1 second of measurement. Trajectories were recorded at a sampling rate of 65 536 Hz.

As the particle position is determined by the deflection of the stationary position detection laser, particle displacement can only be measured correctly in a small area where laser deflection is proportional to bead displacement. In this experiment, the distance is limited to *d*_*max*_ = *±*200 nm. For *D*_*q*_ *>* 0 the particle will quickly leave this detection area, rendering the experiment impractical. To overcome this limitation we generate the trap trajectories prior to the experiment. Iterating through the trajectory, each time a position *x*(*t*_*i*_) = *±d*_*max*_ is reached at a time *t*_*i*_, the left over trajectory is mirrored at this boundary according to

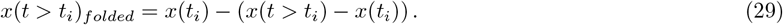

In this manner, the entire trajectory is folded into the area between *−d*_*max*_ and *d*_*max*_ where accurate position detection is possible. After the particle trajectory was measured, it was unfolded in the same manner to recover the intended active motion (Extended Data Fig. 1). Since the motion is generated from a fully random underlying process, and the relaxation timescale is much shorter than the switching timescale, no systematic effect of this approach is to be expected.

### M 5 Active and passive microrheology in living cells

For active and passive microrheology, the previously described optical tweezers setup was used. Cells were prepared according to Material and Methods M 7 and mounted onto the setup. During experiments, cells were kept at 37 *±*C to ensure physiological conditions.

For each cell type or condition at least N=60 cells were investigated on n=3 subsequent days. Data points were recorded at a sampling rate of 65 536 Hz

#### Active microrheology

In active microrheology, the trapping laser is oscillating with a variable driving frequency *f*_*D*_ and thereby applying a sinusoidal force *F* (*t*) onto a probe particle inside of a cell. Simultaneously, the second position detection laser is recording the particle position *x*(*t*). For such a measurement, particle position and force are linked via the response function *χ*(*t*):

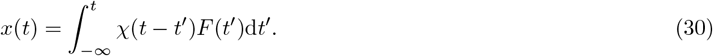

In Fourier space, this convolution can be evaluated as a product which yields access to the response function,

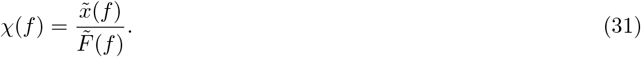

which characterizes the viscoelastic properties of the probed area. To capture the full frequency dependent response function, the probe particle is driven successively at different driving frequencies (*f*_*D*_ ∈ {1Hz, 2Hz, 4Hz, 8Hz, 16Hz, 32Hz, 64Hz, 128Hz, 256Hz, 512Hz, 1024Hz}). For each frequency at least 3 periods where recorded up to a maximum time of 1 second. Using the equilibrium part of the generalized Stokes-Einstein relation^54^

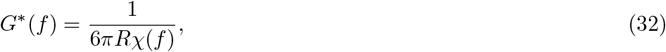

with R being the particle radius, we determine the complex shear modulus *G*\*(*f*).

#### Passive fluctuation measurements

Passive fluctuations measurements have been used to acquire passive microrheology in system with only thermal fluctu-ations. To determine the MBR, a precise access to particle fluctuations by pure passive observation provides the basic measurement. Here we monitor the probe particle motion in absence of any optical tweezers forces. This is done using solely the position detection laser. For each cell, the position signal was recorded 3 times over a period of 10 seconds. This measurement is then used to calculate the mean back relaxation but also to determine the the power spectral density 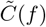 which give access to the activity of the system.

In thermal equilibrium, particle fluctuations in terms of the power spectral density

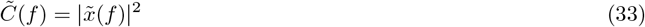

are directly linked to the dissipative part of the response function (measured with active micro rheology (AR)) *χ″*(*f*) via the thermal energy:

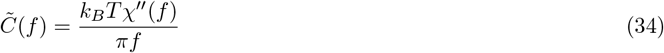

In active systems, like cells, additional metabolic energy is provided to the system. Therefore thermal energy (*k*_*B*_*T*) is not sufficient to describe the particle motion 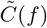. We introduce an effective energy *E*_eff_ which compensates for this additional active energy *E*_*a*_:

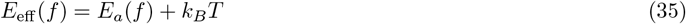

In active systems, the effective energy is then given as

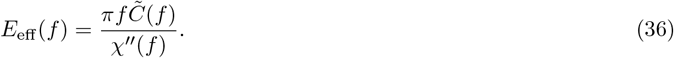

A deviation from 1*k*_*B*_*T* of this effective energy indicates non-equilibrium properties of the system. In this adapted version of the FDT that is extended to non-equilibrium situations, it is possible to determine the dissipative response function if both the power spectral density and the effective energy are known.

### M 6 Data analysis and statistics

#### Motor model

For each level of activity *D*_*q*_, 100 particle trajectories were recorded with each period being 1 second. Subsequently, the MBR was calculated. The presented data shows the mean plus/minus standard deviation.

Effective energies were calculated as the ratio of the mean experimentally determined power spectral density 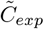 (see SI 3) and the theoretically expected PSD of a passive system 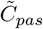 with measured values of stiffness *k* and and friction coefficient *γ*:

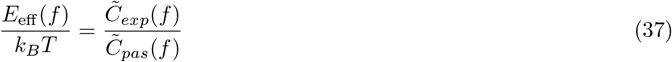

Here we use that in passive systems the fluctuation dissipation theorem holds true: 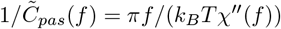.

#### Active cells

For each cell type or condition, N individual cells were investigated in total on n subsequent days: The results shown for effective energy *E*_eff_, response function *χ*, power spectral density 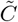 and complex shear modulus *G*\* are given as median and standard error of the mean (SEM) if not stated otherwise.

**Table 1:** Amount of experiments per cell type or condition. N is the total number of investigated cells. n is the number of independent experimental days.

| Cell type | N | n |
| --- | --- | --- |
| A549 | 62 | 3 |
| C2C12 | 57 | 3 |
| CT26 | 64 | 3 |
| HeLa | 79 | 6 |
| HeLa passivated | 25 | 3 |
| HoxB8 | 61 | 4 |
| MCF7 | 61 | 3 |
| MDCK | 65 | 3 |

#### Deriving the complex shear modulus from the long time MBR limit

To get the phenomenological relation between MBR and effective energy amplitude *E*_0_, the MBR longtime value (MBR(1*s*)) of all investigated cell types was plotted against the corresponding amplitude *E*_0_. *E*_0_ was obtained by fitting the median effective energy with the function 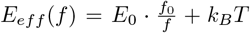. Here *f*_0_ is fixed to 1Hz, to keep the units consistent. A linear fit of this data yields a phenomenological relation between long time MBR limit and effective energy prefactor *E*_0_ (Fig. 4b) for all measured cells:

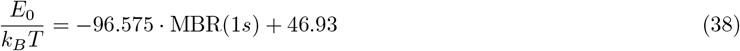

This relation could then be used to derive the effective energy from the MBR of any new cell type. To test this statement we applied it to MCF7 cells, a new cell type that was not used to derive the relation between *E*_0_ and MBR. Using Eq. 38, the effective energy amplitude *E*_0_, and with this also the full effective energy spectrum *E*_eff_ (*f*), was predicted from the purely passive measurement of the MBR. Additionally, the same passive measurement gives access to the power spectral density 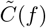. With the knowledge of *E*_eff_ (*f*) and 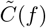, we can exploit Eq. 36 to obtain the imaginary part of the response function *χ″*(*f*). As the imaginary and real part of the response function are not independent, knowledge of the imaginary response function is sufficient to fully obtain the viscoelastic material properties. To recover the real part, either the Kramers Kronig relation can be used, or a model can be fit to the data. As it was shown in a series of recent studies that the complex shear modulus *G*\*(*f*) is well described by a double power law,^3, 23^ we use this model to model the mechanics of the cell.:

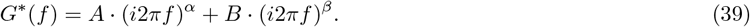

Using the relation *G*\*(*f*) = 1*/*(6*πRχ*(*f*)) we find an analytic expression for *χ″*(*f*) as

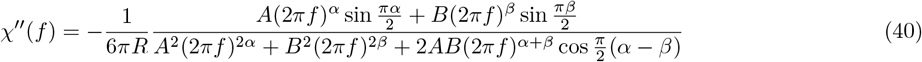

which is taken to fit the parameters *A, α, B* and *β*, that then fully describe the viscoelastic shear modulus *G*\*(*f*). As presented in figure 4 c,d,e, we can now quantify the deviation between *χ″*(*f*), *G′*(*f*) and *G″*(*f*) obtained from established active microrheology (AR) and from the new MBR approach. The relative deviation *R* is defined as:

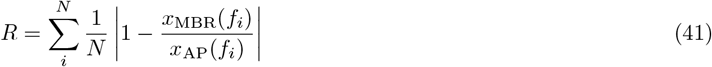

where *x*_MBR_(*f*) is the value of a quantity of interest at frequency f.

### M 7 Cell culture and bead insertion

A549, C2C12, CT26, HeLa and MDCK cells were cultured at 5% CO_2_ and 37°C in Dulbecco’s modified Eagle medium (DMEM, Capricorn) with 1% Penicillin Streptomycin (Gibco) and 10% fetal bovine serum (FBS, Sigma-Aldrich). HoxB8 cells were provided by the Institute for Immunology, University of Münster, Germany and cultured according to.^55^ 3 days before the experiment, HoxB8 cells were suspended in differentiation medium for differentiation into macrophages by the time of experiment. Except for the culture medium and using EDTA instead of Trypsin, the following procedure was the same for HoxB8 cells.

Before experiments, the cells were split close to full confluency and seeded on Fibronectin-coated glass cover slips (22×55×0.15 mm VWR). Previously 1 μm beads (Polybead^®^ Microspheres 1 μm, Polyscience, Inc) were added in 1:10,000 dilution to the medium. After this, cells where incubated for up to 15 hours until a sufficient amount of beads was phagocytozed by the cells. Cells were then washed with PBS in order to remove the left over beads outside the cells. After aspiration of PBS, a smaller cover slip was glued on top of the estimated cell spreading area using 2 layers of 200 μm adhesive tape (DST1950, Thorlabs, New Jersey, USA). The sample chamber was filled with CO_2_ independent medium (CO_2_ Independent Medium, Gibco™) in order to provide a healthy environment for the cells during the experiments.

### M 8 Pharmacological treatment

Cytochalasin B (CB)and Nocodazol (Noc) were purchased from Sigma-Aldrich. Both drugs were used at a concentration of 10 μg mL^*−*1^ and added to the medium 10 minutes prior to the experiment.

## Data availability

All source data can be found at:

https://owncloud.gwdg.de/index.php/s/GQ4KHRj4eHvwwnt

## Code availability

The code used for data analysis can be founde at

https://gitlab.gwdg.de/tillmoritz.muenker/mean-backward-relaxation

## Acknowledgements

We thank the Institute for Immunology from the University of Münster for providing us with HoxB8 cells and the corresponding resources. The authors thank M. Kardar and J. Enderlein for helpful discussion. TM and TB have received funding from the European Research Council (ERC) under the European Union’s Horizon 2020 research and innovation programme (PolarizeMe, Grant agreement No. 771201). The authors thank R. Jain for discussions at the early stages of this work.

## Author information

### Affiliations

**Third Institute of Physics, Georg August Universität Göttingen, Göttingen, Germany**

Till M. Muenker, Timo Betz

**Institute for Theoretical Physics, Georg August Universität Göttingen, Göttingen, Germany**

Gabriel Knotz, Matthias Krüger

## Contributions

T.B and M.K conceived the study. M.K and G.K established the theoretical framework of MBR. T.M.M designed and performed all experiments. T.M.M performed data analysis. G.K performed simulations. All authors performed data interpretation, discussion and wrote the manuscript.

## Extended data figures

**Extended Data Figure 1:**
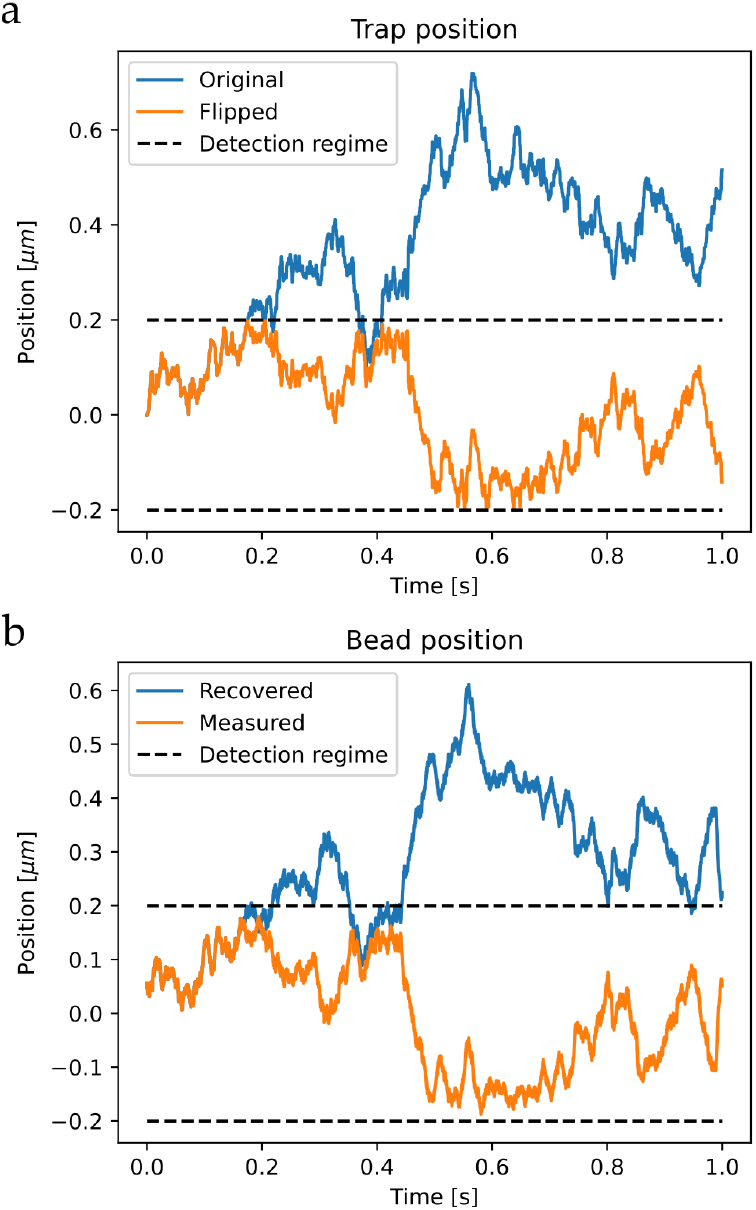
Trajectory flipping. a For certain levels of activity *D*_*q*_, the trap trajectory (blue) would exceed the area were position detection is possible. Therefore the original trajectory was flipped each time it would leave the indicated regime. b The bead position in response to the trap position was measured (blue). To recover the trajectory of a particle subject to a trap diffusion *D*_*q*_, the trajectory was flipped at equal time points to the trap trajectory (orange).

**Extended Data Figure 2:**
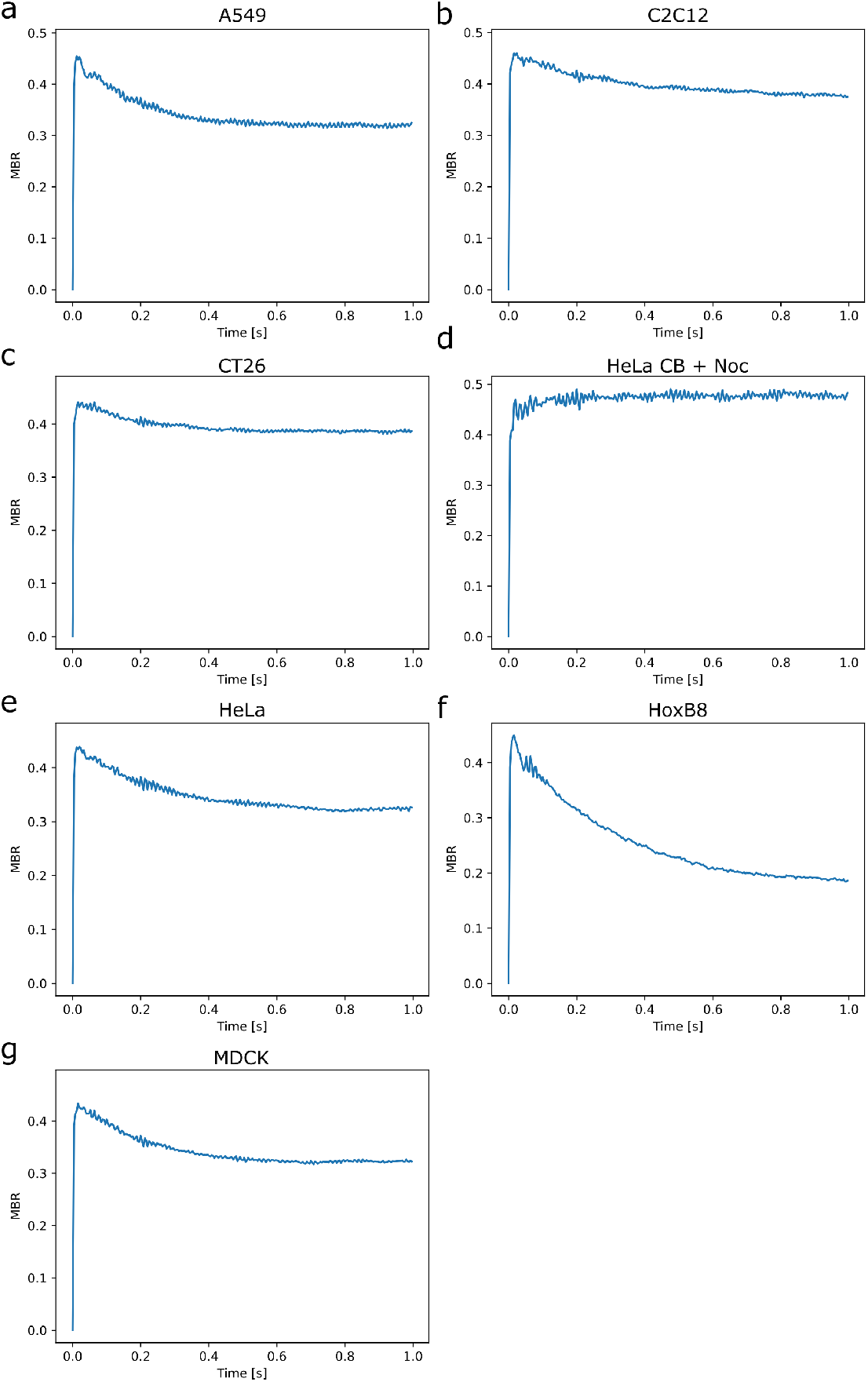
MBR calculated for individual cell types

**Extended Data Figure 3:**
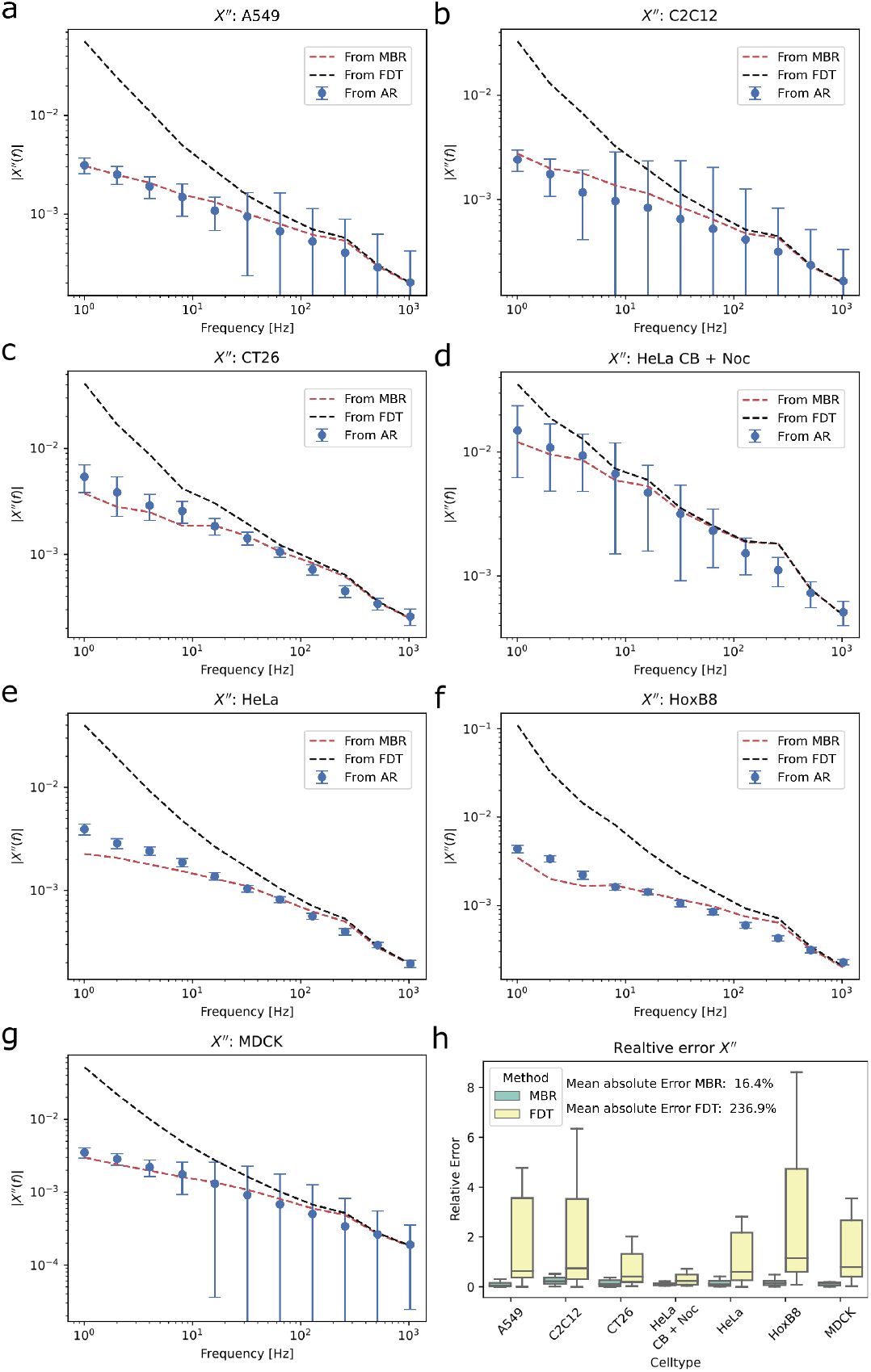
Comparison of MBR approach, classical active and passive rheology approach and naive approach assuming validity of the FDT. a-g Response functions of all cell types derived from active rheology AR, derived assuming a passive system and using fluctuation dissipation theorem FDT and derived using the novel MBR approach. h Relative error between MBR/FDT method and the established AR approach. For all cell types, the novel MBR approach shows less deviation from AR results.

**Extended Data Figure 4:**
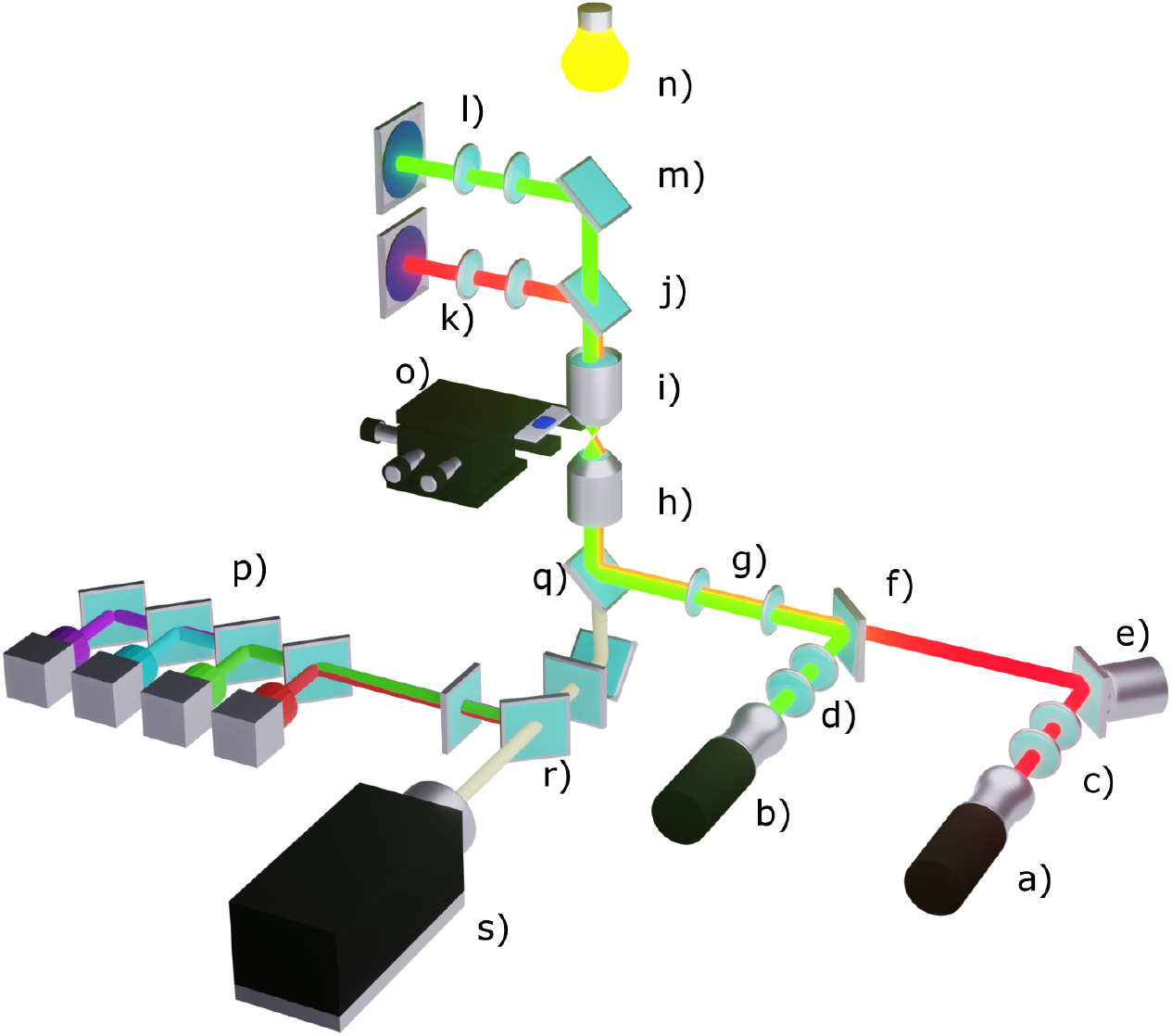
Optical tweezers setup. a) 808 nm trapping laser. b) 967 nm detection laser. c) 5x telescope. d) 3x telescope. e) piezo tilting platform with mirror. f) dichroic mirror. g) 4f configuration. h) 60x NA 1.2 Objective. i) NA1.4 condenser. j) dichroic mirror. k) BFP projection onto PSD. l) BFP projection onto QPD. m) dichroic mirror. n) 620 nm brightfield LED. o) stage. p) 405/490/565/660 nm fluorescence LEDs. q) dichroic mirror. r) 4 band beam splitter. s): camera.

**Extended Data Figure 5:**
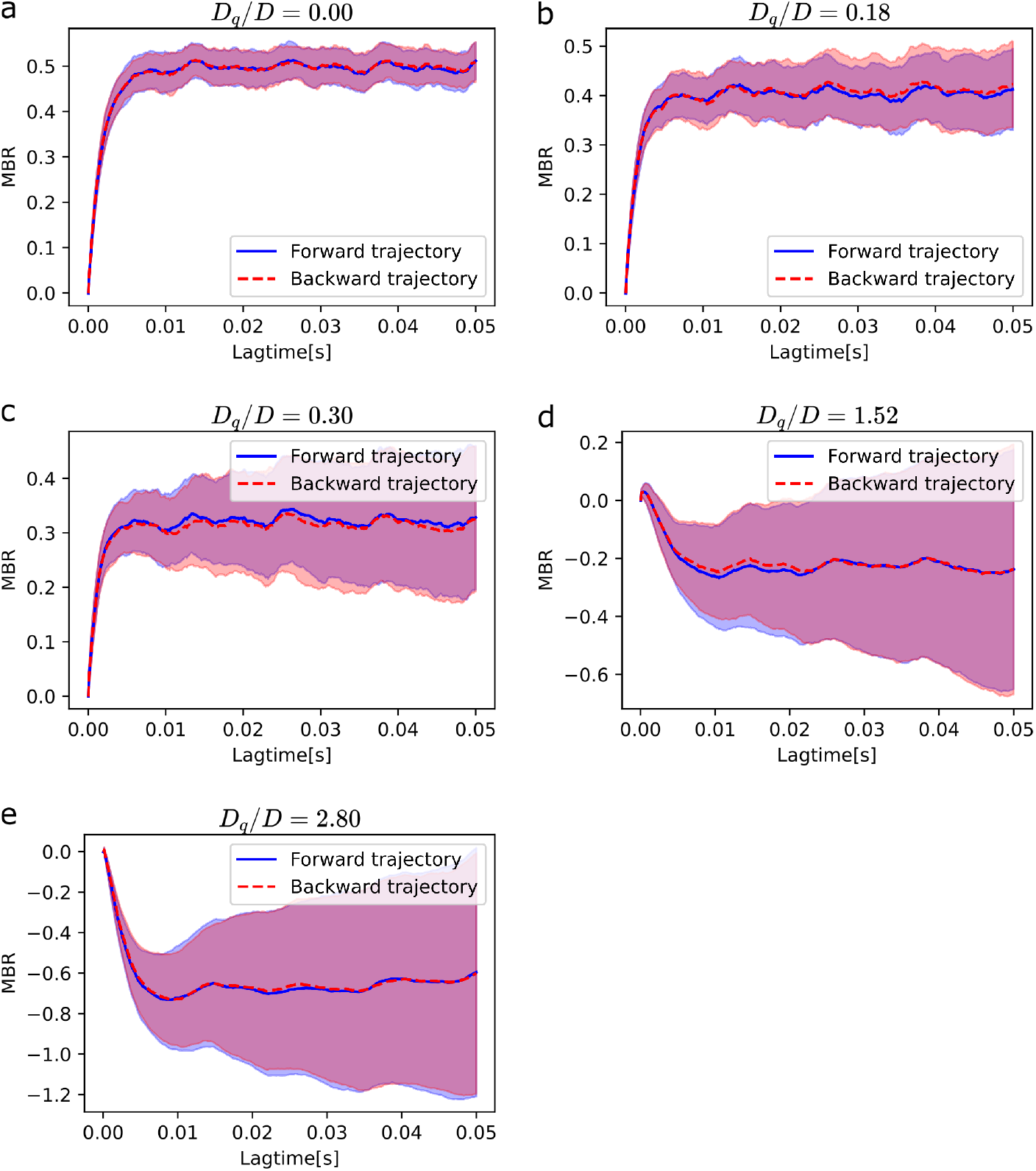
MBR calculated for the motor model. The MBR is determined from original (forward trajectory, blue) and time reversed trajectories (backward trajectory, red). Independent of the level of activity *D*_*q*_*/D*, there seems to be no differences between forward and backward trajectory. Hence, time reversibility is given. Presented errors are given as *±* standard deviation.

## Supplementary Information

### SI 1 Calculation of MBR

Particle trajectories *x*(*t*_*i*_) were typically recorded for a period of 10 seconds with a sample rate of *f*_sample_ = 65.536 Hz. This led to a time resolution of dt = *t*_*i*+1_ *− t*_*i*_ = 1*/f*_sample_ = 15.3 μs.

Formally, the MBR is defined as:

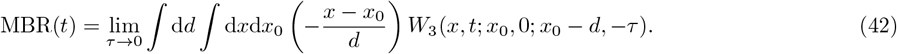

Practically, it calculates the mean distance a particle travels which, previously moved a distance *d* within a short time frame *τ* (see SI 2 for more information about how *τ* was chosen). We allow all kind of *d* and normalize the afterwards moved distance by it. We calculate the distance *d*_*i*_ a particle moved within the time frame *τ* as:

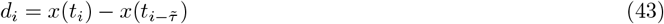

Here 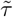 is the lag time *τ* in units of indices : 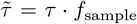. As this displacement *d* will occasionally be smaller than the resolution *d*_*min*_ = 2 ns of the particle tracking system we exclude theses cases by introducing a new index variable *j* which only counts the *N* cases where *d > d*_*min*_. Thus, the MBR is then calculated as:

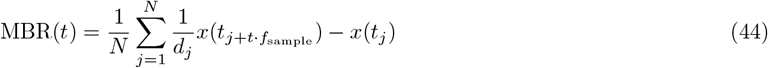

### SI 2 Choosing a value for *τ*

A finite detection accuracy of the probe particle in the experimental setup posse a lower bound to what value can be chosen for *τ*. Even though *τ* should be chosen as small as possible, it should be chosen large enough, such that the probe particle has the chance to move a larger distance than the detection accuracy. The setup used for all experiments has an accuracy of 2 nm. To get an estimate for the timescale a typical particle required to travel a mean distance larger than this accuracy, we calculated the mean squared displacement (MSD) of a typical trajectory. By locating the timescale at which the MSD reached the value of the squared accuracy, we obtained an estimate for the optimal value of *τ* (Fig 6).

For all experiments presented in this study, a value of *τ* = 600 μs was chosen.

**Extended Data Figure 6:**
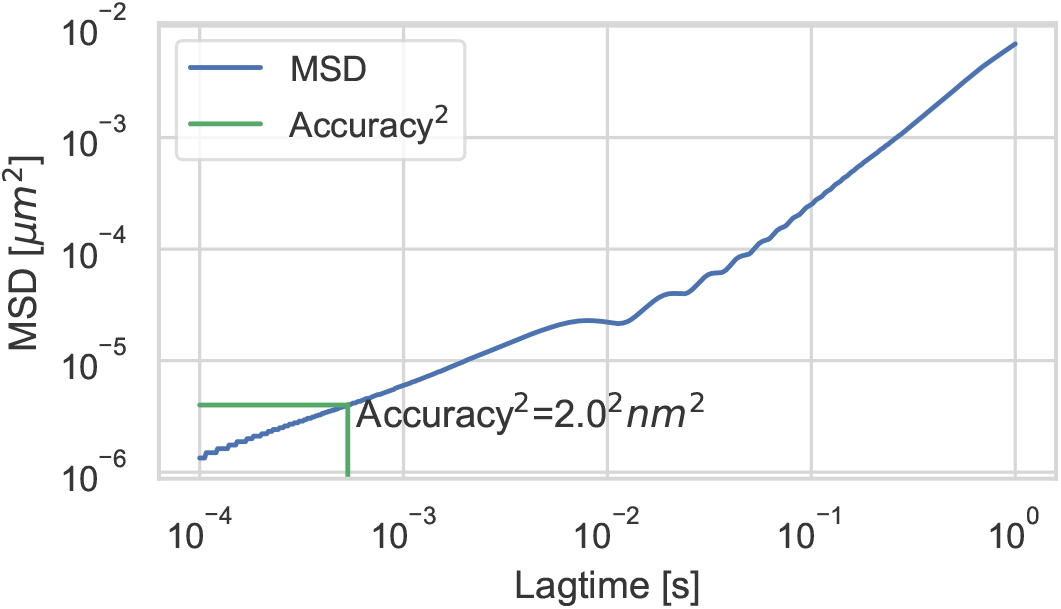
A value for *τ* was chosen by identifying the timescale at which a typical MSD surpassed the squared detection accuracy of 2 nm.

### SI 3 PSD for diffusing potential model

To avoid problems with the Wiener–Khinchin theorem, which connects the correlation function with the PSD, we used the relation between the MSD and the correlation function to get an equivalent measure for the PSD in the motor model. This avoids a possible problem that may arise because the motor model is, mathematically, not in a wide sense stationary process. The MSD and the correlation function are connected via ⟨(*x*(*t*) *− x*(0))^2^⟩ = ⟨*x*^2^⟩ *−* ⟨*x*(*t*)*x*(0) ⟩ where ⟨*x*(*t*)*x*(0) ⟩ is well defined in a stationary process, but not for the motor model. In contrast, the MSD is finite and well defined. With this, we can use the Fourier transform of the MSD as an equivalent measure for the PSD. To numerically perform the transformation we first Fourier transform the second derivative of the MSD and add the transformation of the short time linear term manually.

### SI 4 Effective Energy for diffusing potential

We begin describing the behaviour of a probe particle with size R suspended in a liquid of viscosity *η* trapped in a harmonic potential of stiffness *κ*. The position of the particle is denoted as *x*(*t*) while the position of the trap is denoted with *q*(*t*). The system can be described using 2 stochastic differential aligns.

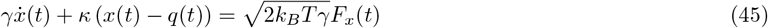

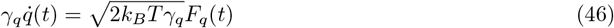

*γ* and *γ*_*q*_ are friction coefficients of the fluid and a pseudo friction for the trap (which does not really feel a friction). *F* (*t*) denotes a stochastic process with properties:

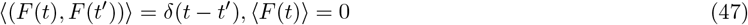

We begin by Fourier transforming the system of differential aligns:

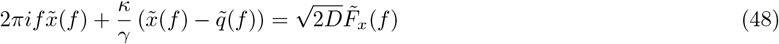

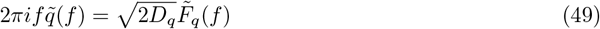

We can now insert 49 into 48 and solve for 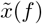:

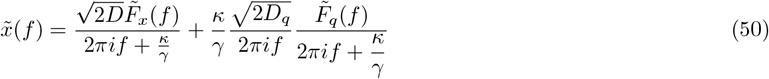

The power spectral density PSD is given as absolute square of the fourier transform: 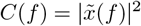. In this calculation the mixed terms drop out and 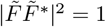 so that we end up with:

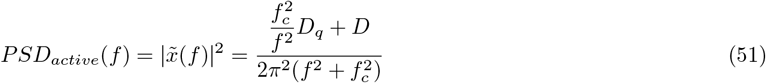

with 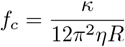 being the corner frequency.

When the trap is stationary (passive system) the dynamics are given by

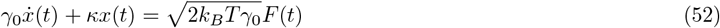

which results in a PSD of :

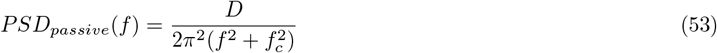

The effective energy in terms of *k*_*B*_*T* is given as the ratio of these power spectral densities:

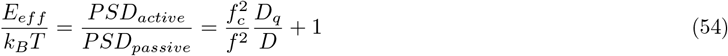

## Footnotes

1 Note that limit of *d →* 0 of Eq. (20) is automatically fulfilled in Eq. (5) in the limit of small *τ*, as the distribution in Eq. (19) gets sharp in *d* due to the term exp(*−d*^2^*/*4*Dτ*).

